# FalseColor-Python: a rapid intensity-leveling and digital-staining package for fluorescence-based slide-free digital pathology

**DOI:** 10.1101/2020.05.03.074955

**Authors:** Robert Serafin, Weisi Xie, Adam K. Glaser, Jonathan T. C Liu

## Abstract

Slide-free digital pathology techniques, including nondestructive 3D microscopy, are gaining interest as alternatives to traditional slide-based histology. In order to facilitate clinical adoption of these fluorescence-based techniques, software methods have been developed to convert grayscale fluorescence images into color images that mimic the appearance of standard absorptive chromogens such as hematoxylin and eosin (H&E). However, these false-coloring algorithms often require manual and iterative adjustment of parameters, with results that can be inconsistent in the presence of intensity nonuniformities within an image and/or between specimens (intra- and inter-specimen variability). Here, we present an open-source (Python-based) rapid intensity-leveling and digital-staining package that is specifically designed to render two-channel fluorescence images (i.e. a fluorescent analog of H&E) to the traditional H&E color space for 2D and 3D microscopy datasets. However, this method can be easily tailored for other false-coloring needs. Our package offers (1) automated and uniform false coloring in spite of uneven staining within a large thick specimen, (2) consistent color-space representations that are robust to variations in staining and imaging conditions between different specimens, and (3) GPU-accelerated data processing to allow these methods to scale to large datasets. We demonstrate this platform by generating H&E-like images from cleared tissues that are fluorescently imaged in 3D with open-top light-sheet (OTLS) microscopy, and quantitatively characterizing the results in comparison to traditional slide-based H&E histology.

## Introduction

Modern microscopy methods enable life scientists and clinicians to visualize complex tissue structures, where recent technological advancements have radically enhanced our understanding of biological processes and disease pathologies. However, clinical diagnostic practices have not taken full advantage of these modern microscopy techniques. In particular, the gold-standard diagnostic method of histology is based on centuries-old technologies, where tissues are preserved in harsh fixatives, destructively sectioned onto glass slides, stained with simple chromogens (most-commonly with hematoxylin and eosin, i.e. H&E), and manually imaged with analog brightfield microscopes. In order to improve throughput, non-destructiveness, sampling extent, and in some cases, to provide 3D information, several slide-free microscopy techniques have recently been explored for use in clinical settings. For example, techniques such as confocal microscopy [1, 2], multiphoton microscopy [3-6], microscopy with UV surface excitation (MUSE) [7-9], and structured illumination microscopy (SIM) [10, 11] have been explored as slide-free alternatives to frozen-section histology for rapid interoperative guidance. Additionally, light-sheet microscopy technologies, such as open-top light-sheet (OTLS) microscopy [12 -14], when used in conjunction with tissue-clearing techniques [15], have been explored for slide-free nondestructive 3D pathology. These techniques generally rely on fluorescence collected with a sensitive monochrome detector. Since pathologists are accustomed to certain color schemes generated by standard chromogens, such as the pink and purple hues associated with H&E staining, the ability to render grayscale fluorescent images with color palettes that mimic standard histology can play a major role on the ability of pathologists to interpret and adopt slide-free pathology methods in the future.

Several groups have published software methods to convert two-channel fluorescence images into “virtual H&E” images. One such virtual H&E algorithm was published in 2009 by Gareau [16]. Using an additive model, this method rendered the reflectance contrast generated by collagen and cytoplasm, and the fluorescence generated by a hematoxylin analog, to mimic H&E staining. Bini et al. in 2011 further refined this model by analyzing the transmitted spectra of multiple slides that were independently stained with either hematoxylin or eosin [17]. A limitation of the additive approach is that in standard H&E histology, the classic pink and purple hues are the result of spectral mixing of the two dyes present in the specimen, according to a nonlinear absorption process (Beer-Lambert-law attenuation). Additive models rely on the linear superposition of intensities, which is non-physical and does not reliably mimic the appearance of conventional H&E histology.

To address these limitations, a false-coloring model based on the Beer-Lambert law of absorption was developed by *Giacomelli et al*. in 2016 [18]. The Beer-Lambert model accounts for the spectral mixing of chromogenic (absorption-based) dyes and their wavelength-dependent nonlinear attenuation of light (i.e. exponential decay as a function of concentration). As a result, this model can accommodate multiple fluorescent stains with overlapping spectra without loss of contrast or non-physical results. Note that in addition to H&E, other chromogens can be modeled, such as the DAB stain most-commonly used for immunohistochemistry (IHC) [13].

The previous approaches described above, for false coloring fluorescent images to mimic conventional chromogenic (absorption-based) stains, have a number of practical shortcomings that prevent them from generating consistent results:

1. Significant variations in the appearance of slide-free digital pathology images can occur due to uneven staining within a specimen (intra-specimen variability). In particular, thick unsectioned tissues exhibit diffusion barriers that often lead to nonuniform staining. This is more pronounced with large agents such as antibodies but also affects smaller agents. Additionally, when imaging optically cleared tissues, samples that are not fully cleared can exhibit a scattering-induced loss of signal as a function of imaging depth.
2. The appearance of both standard histology and slide-free fluorescence images can vary greatly between institutions, or even within an institution, due to day-to-day variations in sample preparation, staining protocols and imaging parameters (inter-specimen variability) [19]. In the presence of these variabilities, simple false-coloring algorithms can fail unless software parameters are manually tweaked, often by trial-and-error.
3. Finally, 3D slide-free digital pathology datasets are often hundreds of gigabytes to terabytes in size, which is 3 to 4 orders of magnitude larger than standard 2D whole slide images. To facilitate clinical translation, false-coloring algorithms must be able to process these large 3D datasets in an efficient and scalable manner.

It bears mentioning that several deep-learning methods have been developed to render H&E-like images from unstained tissue sections imaged with label-free imaging modalities such as brightfield and autofluorescence microscopy [20-22], multi-photon microscopy [23], and quantitative phase-contrast imaging [24]. Similar machine-learning-based approaches could be used to create H&E-like images from tissues labeled with exogenous fluorophores (i.e. fluorescent analogs of H&E). However, these methods not only require large amounts of training data but are also highly sensitive to pre-analytical variations such as staining protocols, imaging parameters, and hardware settings. In contrast, the ability to use an explainable physics-based approach for digital staining is likely attractive for many end users and regulatory agencies, allowing for easier error identification, debugging, and compatibility with different imaging platforms [25].

In this manuscript we present FalseColor-Python, which enables rapid and robust digital staining of fluorescence images to mimic the appearance of chromogenic stains using the Beer-Lambert model. As a pre-processing step prior to false-coloring, an intensity-leveling routine, analogous to flat fielding, is used to locally and globally normalize image intensities, thereby mitigating the effects of both intra- and inter-sample variability. In particular, the code presented here is specifically optimized to generate H&E-like images of tissues stained with a two-channel fluorescent analog of H&E (i.e. a fluorescent nuclear stain that mimics hematoxylin, along with the stromal stain, eosin, which is naturally fluorescent) and imaged with 3D open-top light-sheet (OTLS) microscopy [12-14]. We show that FalseColor-Python enables accurate and reproducible H&E false-coloring that qualitatively and quantitatively matches the appearance of H&E whole slide images, but with less variability than is seen with standard histology. GPU acceleration is used for efficient and scalable processing of large 3D datasets.

Note that while H&E false-coloring is demonstrated here, FalseColor-Python can be tailored to accommodate other preferred fluorescent staining combinations and individual color preferences. For example, FalseColor-Python can be adapted to false-color fluorescence images to mimic other chromogens, such as DAB (used in standard immunohistochemistry [13]) or other special stains (e.g. PAS, Masson’s Trichrome, Toluidine Blue). In addition, while FalseColor-Python is optimized for large 3D datasets, it can easily be applied to 2D images as well.

### FalseColor-Python workflow

Figure 1 outlines the workflow for processing large two-channel 3D datasets using FalseColor-Python. For our specific implementation of this code, input data is stored on disk in the Hierarchical Data Format (HDF5), which is a common multi-resolution 3D image format (**Fig 1A**). The advantage of multi-resolution file formats like HDF5, or similar formats such as N5, is that they contain multiple down-sampled versions of the imaging data. For our 3D microscopy data, we use a 16x down-sampled version of the data (low-resolution) to rapidly calculate an approximate global background level for each channel. The background level is calculated by first ignoring all pixels that are below a threshold value (i.e. black pixels), which we have defined as 4 standard deviations above the detector noise floor. The intensity of the 20^th^ percentile of the remaining pixels is defined as the background value, which is due to a combination of tissue autofluorescence and nonspecific staining. We have found that for all of the 3D datasets we have examined, this 20% threshold is effective. However, for tissues that are very densely stained, in which there are few “background” pixels, this threshold may need to be adjusted. Once calculated, this background level is uniformly subtracted from both the down-sampled and full-resolution original dataset (**Fig 1B**). Next, the down-sampled dataset is partitioned into uniformly sized data cubes. In our case, each data cubes corresponds to a 100 × 100 x 100 µm^3^ volume of tissue (where 100 µm corresponds to 256 pixels in our raw datasets) (**Fig 1C**). We use this partitioned data to create a low-resolution 3D “intensity-leveling map,” which will subsequently be used to remove global intensity fluctuations from the original dataset (see introduction). The intensity-leveling map is generated by calculating the median pixel value for every local data cube (**Fig 1D**), scaling it with an empirical weighting factor, *α*, and then linearly interpolating between the cubes to smooth out the map (**Fig 1E**). Intensity leveling is achieved by taking a pixel-by-pixel ratio of the full-resolution dataset (background-subtracted) and the interpolated intensity-leveling map (**Fig 1F**). The result is an image in which the median value of every local region (i.e. data cube) is approximately equal (due to the ratioing step), in which many of the large-scale (gradual) intensity irregularities are reduced (**Fig 1G**). Finally, the leveled images from both channels are passed into the Beer-Lambert false-coloring algorithm, which generates a color (RGB) image (**Fig 1H**). The intensity-leveling and virtual H&E routines are accelerated with a GPU using the CUDA framework from Python’s Numba library [26].

**Figure 1.**
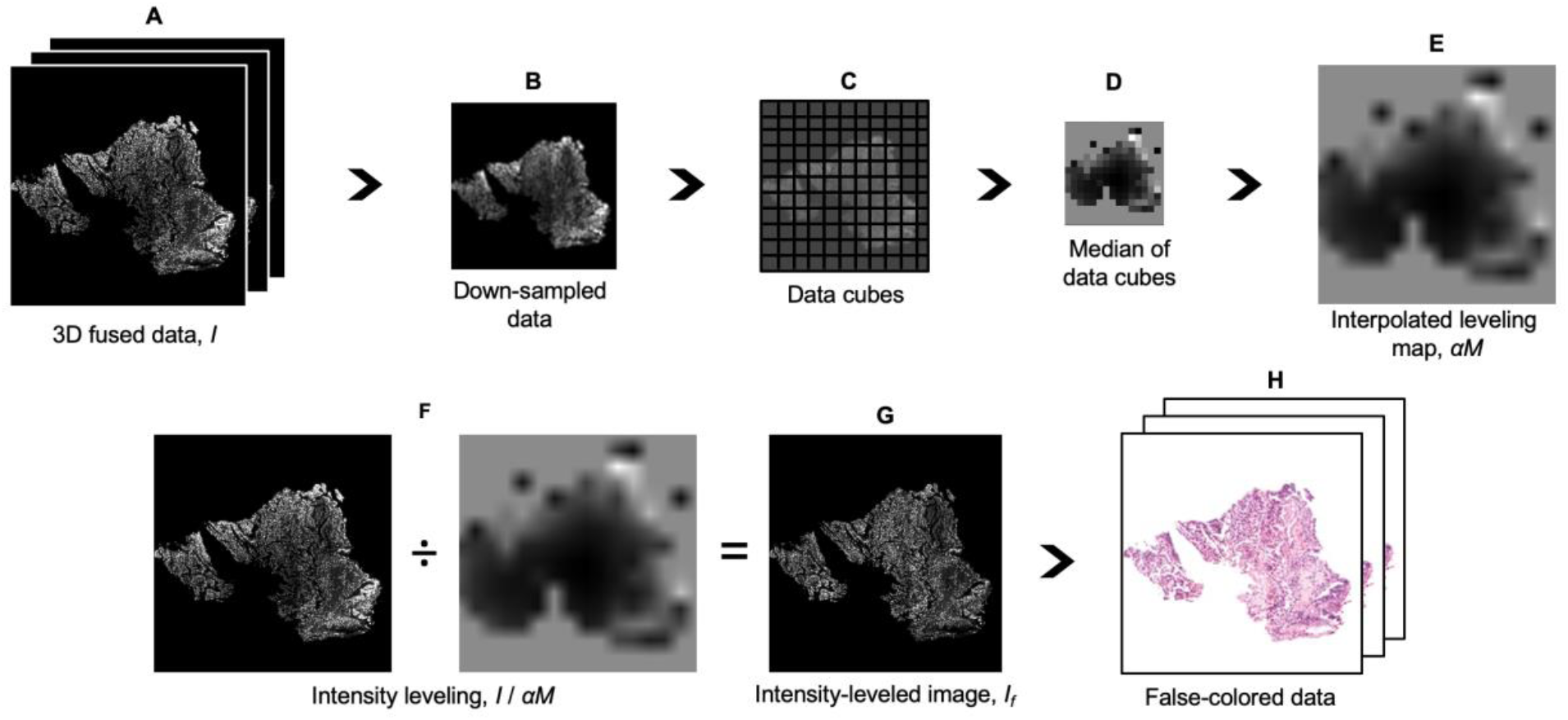
FalseColor-Python workflow for virtual H&E rendering of a two-channel fluorescent analog of H&E. The following operations are performed on both channels, but only one is shown for simplicity. **(A)** 3D data (*I*) is loaded from disk. **(B)** A down-sampled version of the dataset (16x down-sampled here) is extracted, and a background level is calculated for each channel, which is uniformly subtracted from both the down-sampled and full-resolution original dataset. **(C)** The down-sampled data is further subdivided into cubes. **(D)** A preliminary 3D leveling map is generated by calculating the median pixel value for each data cube. **(E)** A full-resolution leveling map, *αM*, is then generated by interpolation. Here, *α* is an empirically determined constant that controls image brightness. **(F)** To achieve intensity leveling, the full-resolution data, *I*, is divided by *αM*. This evens out coarse non-uniformities within the image. **(G)** The leveled images, *If* for both channels are input into the Beer-Lambert model to generate virtual H&E images, **(H)**.

Note that the size of the data cubes (**Fig 1C**) that are used for generating an intensity-leveling map should be chosen based on the spatial scale of the intensity nonuniformities that one wishes to correct. For images that exhibit very gradual or minimal intensity variations, larger cube sizes can be used. Small cube sizes will remove finer-scale intensity fluctuations but may also remove fluctuations/contrast that are due to real tissue structure rather than staining artifacts.

To demonstrate the utility of this method, a comparison of virtual H&E staining with and without intensity-leveling is shown in **Fig 2**. Here a single plane from a 3D microscopy dataset of an optically cleared lung specimen is shown. The specimen was stained with a fluorescent nuclear stain TOPRO-3 Iodide (Cat: T3605, Thermo-Fisher) and was optically cleared with Ethyl-Cinnamate (Cat: 112372, Sigma-Aldrich) [14]. The cleared tissue volume was then imaged using OTLS microscopy [27]. The Beer-Lambert false-coloring algorithm was applied to a single grayscale image without intensity leveling (**Fig 2A**). In this example, the exterior of the tissue was stained more heavily than the interior. In **Fig 2B**, the intensity-leveling procedure was incorporated into the false-coloring routine, such that the false-colored nuclei appear much more uniform across the entire image.

**Figure 2.**
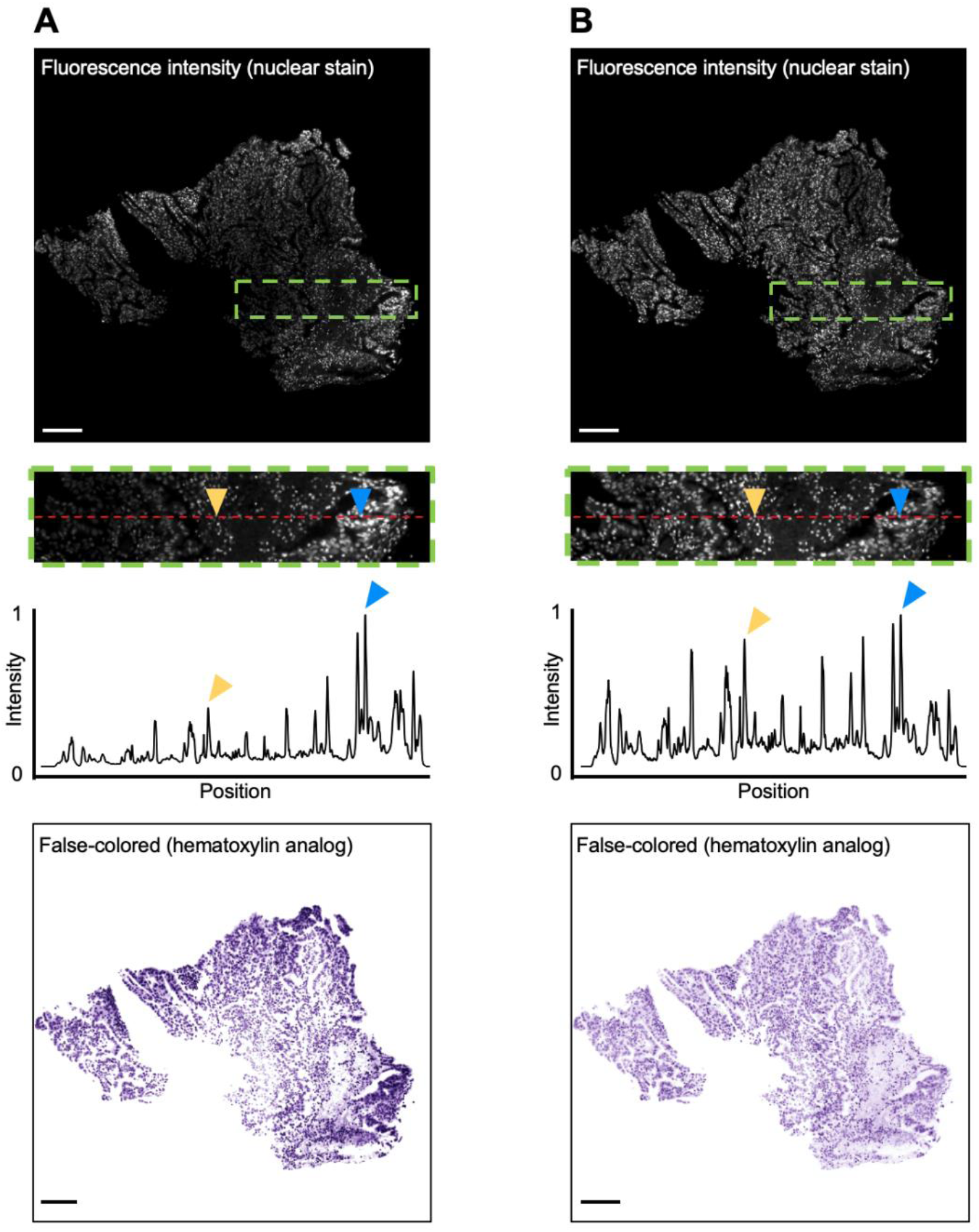
False-coloring with and without intensity-leveling. Only one channel is shown for simplicity. **(A)** Images are shown of a thick tissue specimen that is labeled (nonuniformly) with a fluorescent nuclear stain (TO-PRO-3), optically cleared, and then imaged with 3D microscopy. The image is false colored to mimic hematoxylin without intensity-leveling applied. A line profile within the inset clearly shows that nuclei in the inner region of the tissue (yellow arrow) are dimmer than nuclei on the exterior (blue arrow). When false colored, the nuclei on the exterior appear much darker (bottom panel). **(B)** The same field of view is shown after intensity leveling is applied. Nuclei across the field of view exhibit similar intensities, and when false colored, are much more uniform in appearance. Scale bars: 100 µm

### Colorimetry to mimic standard histology

For the clinical adoption of slide-free digital pathology, virtual H&E algorithms must consistently render virtual H&E images that qualitatively and quantitatively match the coloration of standard histology. To ensure that FalseColor-Python’s virtual H&E algorithm mimics the color palettes of standard histology, we measured the color properties of 65 publicly available whole slide images (H&E) of prostate adenocarcinoma from the Cancer Digital Slide Archive [28] (**Fig 3A**). 10 regions of interest (ROIs) were selected from each whole slide image at a magnification of 200x (20x objective), **(Fig 3 B**). Hematoxylin and eosin stains were segmented from each region of interest using color deconvolution [29, 30]. The color deconvolution algorithm generated a probability map for each dye, where pixel values represent the probability of that pixel belonging to one of the top-two color components of the image (i.e. hematoxylin or eosin), (**Fig 3 C & D**). These probability maps were used to create binary masks for each structure using Otsu’s thresholding method [31]. The binary masks yielded segmented images of the nuclei (hematoxylin stain) and stroma (eosin stain). This process was repeated for all ROIs and the median color properties of each stain were quantified and plotted based on the hue, saturation, value (HSV) color model (**Fig 3 E & F**). We used these measured HSV values as a target for our virtual H&E images. In other words, in FalseColor-Python, we have adjusted the color parameters used in the Beer-Lambert false-coloring algorithm so that the median HSV color properties of the virtual H&E images approximate that of standard whole slide images.

**Figure 3.**
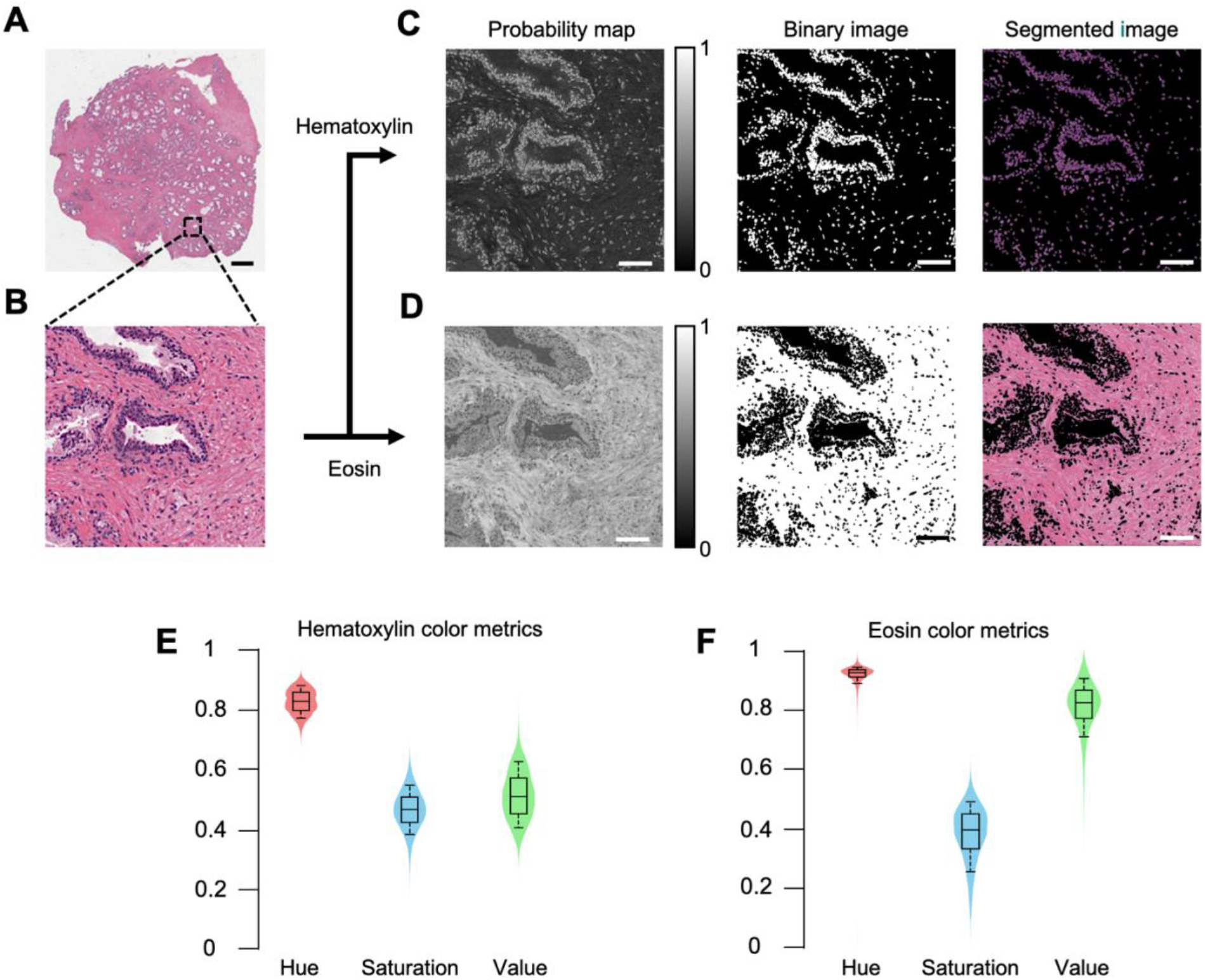
Measuring the color properties of histology images. **(A)** A whole slide image of prostate adenocarcinoma from the Cancer Digital Slide Archive, *cancer.digitalslidearchive.org*. Scale bar: 2 mm **(B)** Magnified inset of (A). Scale bar: 75 µm **(C, D)** Process to segment the hematoxylin and eosin components. Using color deconvolution, a probability map is generated for the top two components of each ROI from the whole slide image (right panel). A binary mask is created for each of the two components by applying Otsu’s thresholding to the probability map (middle panel). This yields a segmented image of the nuclei (hematoxylin stain) and cytoplasm (eosin stain) (left panel). Scale bar: 75 µm **(E, F)** After segmentation of all ROIs, the median values of hue, saturation, and value (HSV color model) are quantified and plotted for both the hematoxylin-stained and eosin-stained tissue components.

For the clinical adoption of slide-free pathology, virtual H&E images not only need to mimic the color palette of standard H&E but must also be consistent and robust in the presence of variations between samples, for example due to differences in staining protocols or imaging parameters. By incorporating intensity-leveling, our virtual H&E algorithm rendered images with similar appearance despite inter-sample intensity differences. As an example, fluorescent images of two prostate tissue samples are shown in **Fig 4A**, in which significant differences in fluorescence intensities are seen (when imaged with the same device and settings). The intensity levels for each fluorescence channel (TO-PRO-3 and eosin) are shown in the accompanying histograms. Despite these differences in intensity, the resulting virtual H&E images of each sample are qualitatively and quantitatively similar. On the right-most column of **Fig 4A**, a histogram of the value (V) component of each virtual H&E image is plotted and is comparable for both images after the intensity-leveled false-coloring routine is applied.

**Figure 4.**
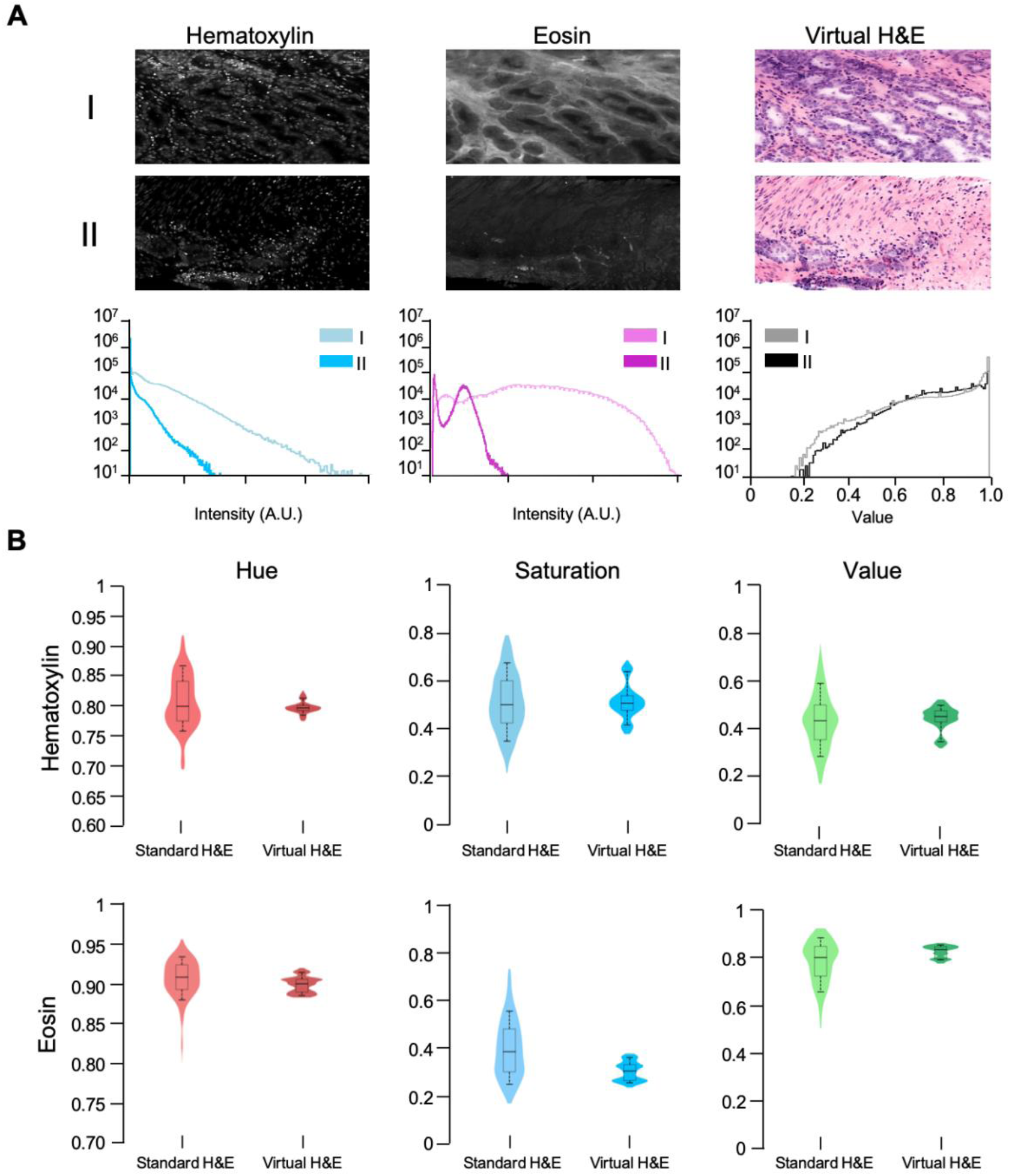
Inter-specimen consistency of virtual H&E images. **(A)** OTLS images of prostate tissue with different staining intensities. Despite significant differences in intensity between example I and II, the virtual H&E images appear qualitatively and quantitatively similar. Histograms are shown of the fluorescence intensities within each image, and the value (V) component of the virtual H&E images. **(B)** Distribution of median color properties for 650 standard H&E and 2100 virtual H&E images of prostate tissue. The distributions for standard H&E exhibit the high degree of color variation seen in conventional histology. The virtual H&E images rendered with FalseColor-Python accurately mimic the coloration of standard H&E with significantly less variation in color parameters than conventional H&E.

To quantitatively compare the consistency of FalseColor-Python with standard H&E histology, we measured the HSV color model parameters for 2100 virtual H&E images from 14 samples of prostate tissue imaged with 3D OTLS microscopy and then processed with FalseColor-Python. To measure the color properties of the virtual H&E images, we used the same color-deconvolution method that was used to measure the color properties of whole slide images from the Cancer Digital Slide Archive (described previously). It should be noted that while the color parameters were adjusted to allow our virtual H&E images to match the color parameters of standard H&E images, all virtual H&E images used in this analysis were processed in a fully automated fashion using the exact same code parameters for intensity-leveling and coloration. Our results show that the color properties of virtual H&E data processed with FalseColor-Python match the measured values of standard H&E, as expected (**Fig 4B**). Furthermore, the color properties of virtual H&E images processed with FalseColor-Python are much more consistent (less standard deviation) than the color properties of standard H&E-stained whole slide images. **Table 1** lists the median and standard deviation of each color property for both standard and virtual H&E. Finally, an image atlas is shown in **Fig 5** to demonstrate that FalseColor-Python renders consistent virtual H&E images across tissue types and to demonstrate that FalseColor-Python is easily adjusted to mimic other color spaces in standard histology (DAB staining).

**Table 1:** Measured median color properties of standard and virtual H&E images in HSV color space. Standard H&E properties are measured from 10 ROIs from each of 65 whole slide images (prostate). Virtual H&E properties are measured from 2100 two-dimensional “optical sections” from 14 prostate specimens.

|  | *Hematoxylin* |  | *Eosin* |  |
| --- | --- | --- | --- | --- |
|  | *Standard* | *Virtual* | *Standard* | *Virtual* |
| *Hue* | $0.811 \pm 0.043$ | $0.808 \pm 0.009$ | $0.910 \pm 0.023$ | $0.901 \pm 0.010$ |
| *Saturation* | $0.453 \pm 0.095$ | $0.457 \pm 0.055$ | $0.381 \pm 0.119$ | $0.310 \pm 0.039$ |
| *Value* | $0.494 \pm 0.111$ | $0.502 \pm 0.048$ | $0.788 \pm 0.087$ | $0.811 \pm 0.025$ |

**Figure 5.**
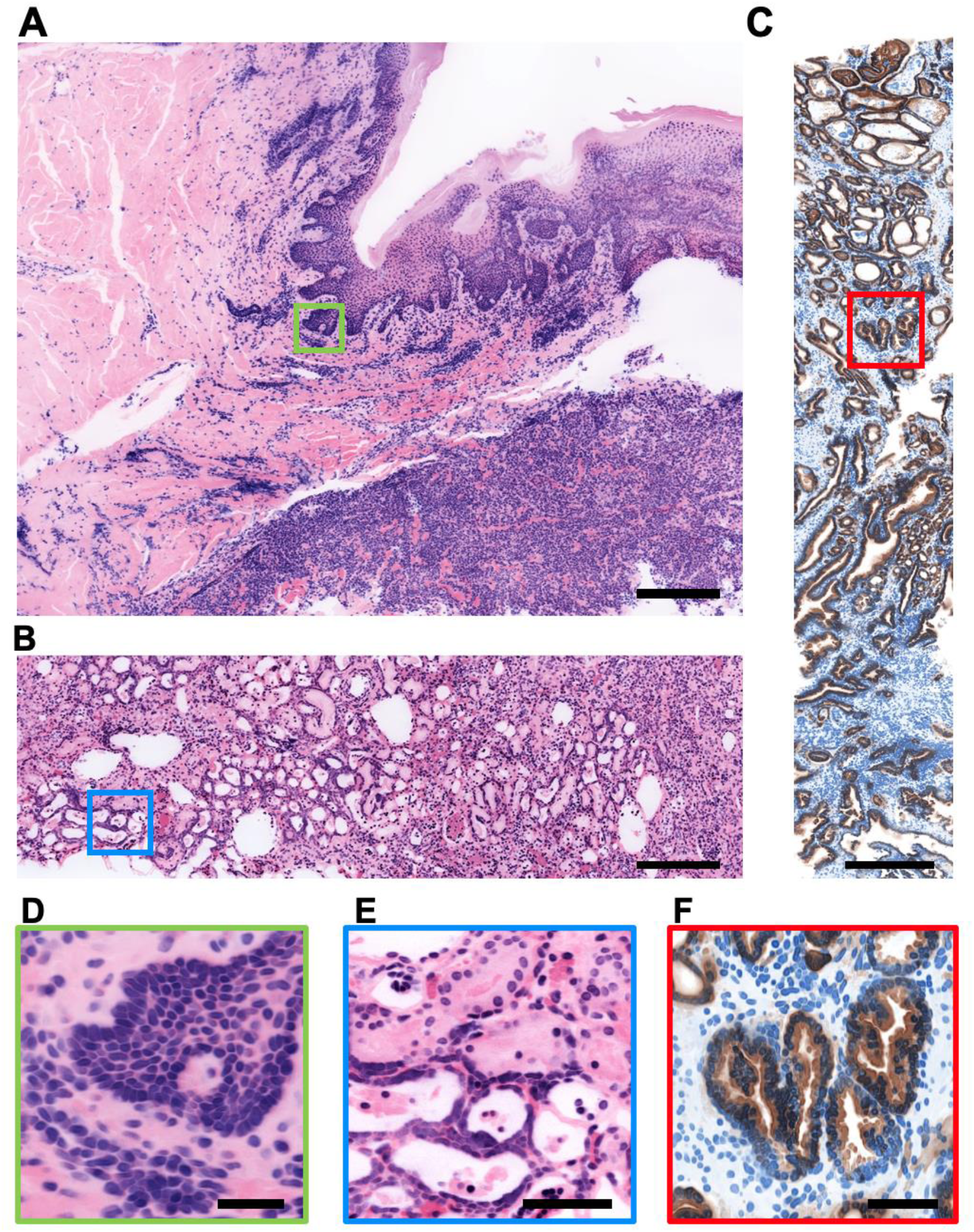
False-coloring image atlas. Thick tissues were stained with a fluorescent analog of H&E **(A, B, D, E)**, or an antibody targeting the high molecular weight keratin (HMWK), CK-8, along with the nuclear stain, TO-PRO-3 **(C, F)**. All tissues were optically cleared and imaged in 3D with an open-top light-sheet (OTLS) microscope. **(A)** Skin. **(B)** Kidney. **(C)** Prostate. **(D)** Basal layer of the epidermis. **(E)** Kidney tubules. **(F)** Prostate glands (carcinoma). Virtual H&E images were processed with identical code parameters in a fully automated fashion. Virtual IHC images were processed with an identical code, but with coloring parameters changed to mimic the chromogen DAB (brown stain) and hematoxylin (blue stain). These codes and parameters are provided in our GitHub repository (see provided link). Scale bars: 500 µm for A-C, 50 µm for D-F.

## Discussion

We have developed FalseColor-Python, a rapid intensity-leveling and digital-staining package for converting grayscale fluorescence images into color images that mimic conventional chromogenic (absorption-based) stains. In particular, we demonstrate the rendering of virtual H&E images from thick tissues stained with a fluorescent analog of H&E and imaged in 3D. To improve the consistency of our false-coloring method in the presence of both inter-sample and intra-sample variations in staining/intensity, we have developed and incorporated a 3D intensity-leveling routine (**Figs 1 & 2**). We analyzed the color properties of standard H&E images (**Fig 3**) and used this data to ensure that the virtual H&E images rendered by FalseColor-Python are representative of standard histology. Our results show that the virtual H&E images rendered by FalseColor-Python are qualitatively and quantitatively similar to standard H&E histology regardless of variations in intensity, as for example due to differences in sample preparation, imaging device, and/or imaging parameters (**Fig 4**). In particular, we have shown that FalseColor-Python renders virtual H&E images that not only quantitatively match the appearance of standard H&E images, but with less variability in coloration than is seen with standard histology (**Fig 4B**).

The ability to render quantitatively and qualitatively consistent virtual H&E images is of critical importance for the adoption of fluorescence-based imaging methods in anatomic pathology. Staining non-uniformities and depth-dependent intensity variations are common data-quality issues in fluorescence imaging, particularly for nondestructive slide-free 3D imaging modalities. In developing this intensity-leveling technique, we have taken advantage of the multiple down-sampled versions of an imaging dataset that are stored in a multi-resolution image format such as HDF5. However, down-sampling of imaging data is straightforward and intensity-leveling maps can be generated from any 3D imaging data. For our 3D microscopy data, optimal performance has been seen with data cubes of 100 × 100 x 100 µm^3^, however users can make adjustments as needed to best suit their data. This tunability is demonstrated in an example located on our GitHub repository. Finally, for certain images with intensity nonuniformities, enhanced performance is seen using an implementation of contrast-limited adaptive histogram equalization (CLAHE) as a pre-processing step before intensity-leveling is performed. This is included as an optional method within FalseColor-Python but was not used in the examples shown in this study.

In terms of limitations, our intensity-leveling method is best suited for images where staining is present throughout the entirety of the sample, but where the spatial variations in that signal are relatively gradual compared with the real high-resolution features of interest (e.g. **Fig 2**.) In cases in which large-scale (gradual) intensity variations are biologically real and informative to end users (e.g. pathologists), there is the possibility that our intensity-leveling methods will mask (i.e. flatten) such global-intensity variations. However, this could be mitigated by tuning the size of the data-partitioning cubes as outlined in **Fig 1**.

A significant outcome of this study is that virtual H&E images rendered by FalseColor-Python exhibit significantly less variability in color parameters than standard histology. This is not surprising considering the obvious differences in appearance seen in histology images generated by independent labs, and even within individual labs at different times. The measured median color-property values of our virtual H&E images were all well within one standard deviation of the color properties of standard H&E images.

We recognize that the standard H&E data used in this analysis represents a subset of all possible color presentations found in histology. Further, we acknowledge that there is a high degree of subjectivity and personal preference regarding the optimal coloration for standard H&E images. Therefore, it is possible to tune the appearance of false-colored images if so desired. For simplicity, RGB color parameters, which are easily measured from any image, are included as arguments in FalseColor-Python’s Beer-Lambert false-coloring algorithm so that users can make adjustments as needed. Further discussion of adjusting color settings is provided in an example on the online repository (see GitHub link at the end of this article).

Note that all methods used for colorimetry analysis of standard and virtual H&E images (e.g. **Fig 3**) are included in FalseColor-Python. Additionally, as mentioned in the introduction, FalseColor-Python is easily adapted for other fluorescence-to-chromogenic staining transformations, for example to render images that mimic chromogenic immunohistochemistry (i.e. DAB stain) (**Fig 5C & 5F**).

FalseColor-Python contains several methods that use GPU acceleration. For example, CUDA-based implementations of the Beer-Lambert false-coloring algorithm are included in FalseColor-Python as well as several preprocessing steps such as intensity-leveling, image sharpening, and background subtraction. For users without access to GPU hardware, equivalent CPU-based processing methods are available as alternatives in FalseColor-Python. In a simple speed comparison, we measured the average time to process 200 two-channel 16-bit images (2048 × 2048 pixels each) with FalseColor-Python, using either GPU- or CPU-based Beer-Lambert false-coloring algorithms and preprocessing steps (i.e background subtraction, intensity-leveling). The GPU-based process was faster by over a factor of 6 (116 +/- 17 ms per image vs. 789 +/- 18 ms per image). There is room for further improvement by developing objects and methods within FalseColor-Python that can process large 3D datasets in parallel. For example, implementing multi-resolution file formats that allow for parallelized read operations (e.g. N5 instead of HDF5) will enable FalseColor-Python to process 3D datasets even more efficiently. Parallelized processing in FalseColor-Python is discussed in an example on the GitHub repository. Developers who are interested in contributing to FalseColor-Python should submit pull requests via the GitHub repository.

## Detailed methods

### OTLS imaging

Tissue samples were stained with a fluorescent nuclear stain, TO-PRO-3 (Cat: T3605, Thermo-Fisher) at a 1:2000 dilution and eosin (Cat: 3801615, Leica Biosystems) at a 1:2000 dilution for 4 hours at room temperature with light shaking. Samples were optically cleared with ethyl-cinnamate (Cat: 112372, Sigma-Aldrich). Stained and cleared samples were imaged on a custom OTLS system [14]. A 660-nm laser was used to excite the nuclear dye, TO-PRO3, and a 561-nm laser was used to excite eosin. Each channel was imaged separately, in succession, with a 16-bit sCMOS camera. For more information on OTLS imaging see our previous publications [12-14].

### Image processing

Two channel OTLS datasets were stored on disk in the HDF5 format with metadata in an XML file structured for analysis using BigStitcher [27]. A custom compression filter (B3D) was used to provide 10x compression. The fine alignment of all OTLS data was performed in BigStitcher and fused to disk as a separate HDF5 file. Before virtual H&E rendering (as described in the manuscript), an optional sharpening routine was used on each image to enhance edges.

Virtual H&E rendering of two-channel fluorescence images was achieved using the Beer-Lambert false-coloring algorithm [18]. Grayscale intensities are converted into RGB images via:

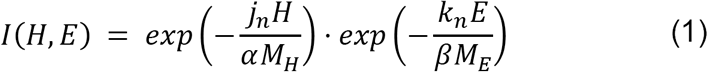

where H, E are fluorescent images, M^H, E^ is the 2D image leveling map for each channel, α, β are intensity-leveling constants, j, k are the color settings and n = R, G, B. Based on the colorimetry measurements described in the manuscript (**Fig. 3**), we adjusted FalseColor-Python’s hematoxylin and eosin color parameters until the measured color properties of virtual H&E images consistently matched those of standard H&E. After this calibration no further adjustments were necessary. **Table 2** lists the RGB color settings used for processing OTLS images. For leveling our OTLS images, *α* = 1.5 and *β* = 3.7 were chosen as scaling constants for the majority of our datasets.

**Table 2:** Reference k values for virtual H&E coloration in RGB space.

|  | *Hematoxylin* | *Eosin* |
| --- | --- | --- |
| *R* | 0.17 | 0.05 |
| *G* | 0.27 | 1.00 |
| *B* | 0.105 | 0.54 |

GPU acceleration for image processing was done using the cuda.jit decorator from the Numba library [26]. Implementation of GPU acceleration is done in such a way that the user needs no experience with the CUDA framework to accelerate their code (see examples on the GitHub repository). To achieve GPU acceleration, users only require a virtual environment equipped with the Anaconda’s cudatoolkit [32] and a CUDA capable GPU. GPU processing was done with a Nvidia Quadro P4000. A script containing the workflow as described in **Fig 1** can be found on the GitHub repository.

### Histology Data Collection

To quantify the HSV color space of traditional histopathology, 65 whole slide images of prostate adenocarcinoma biopsies from the Cancer Genome Atlas, *cancer.digitalslidearchive.org* [28] were examined, 10 equally sized fields of view from each whole slide image were taken at 20x magnification.

### Color-Space quantification

To accurately analyze the color space of histology and OTLS images, the hematoxylin and eosin channels were separated from one another via a binary mask generated by color deconvolution from the *scikit-image.color* python package [30]. This color deconvolution-based segmentation was used for both standard and virtual H&E images. A binary mask was generated for each structure by first applying a median filter to the result of the color deconvolution, and then applying Otsu’s thresholding method [31]. Small objects were removed from the initial mask using an area threshold. A binary opening with a structuring element of a disk, r = 3, was applied to the resulting image. The input RGB image was converted to HSV space and then each mask was applied to generate the final segmented image for each structure. This process was repeated across 650 regions of interest taken from publicly available prostate adenocarcinoma whole slide images and 2100 OTLS virtual H&E images from 14 prostate samples. Once each structure was segmented, the median value for each color property (HSV) was recorded from each segmented image. Zero valued areas, which resulted from the application of the binary mask, were ignored.

### Code Availability

The full FalseColor-python code, example data, and colorimetry analysis methods, are publicly available at *https://github.com/serrob23/falsecolor*. The repository includes instructions for installation and annotated examples.

### Programming Language

Python 3.6+ is used in FalseColor-Python. Some features may be unavailable on older versions. A full list of dependencies and requirements is available on GitHub.

## Acknowledgements

The authors would like to thank Drs. Suzanne Dintzis, Nicholas P. Reder, and Lawrence D. True for assistance regarding histology coloration as well as Lindsey Barner and Kevin Bishop for helpful technical discussions.

## Author contributions

**Software development:** RS, AG.

**Data collection and analysis:** RS, AG, WX, JTCL.

**Wrote the paper:** RS, AG, JTCL.

## References

1. Longo C, Ragazzi M, Rajadhyaksha M, Nehal K, Bennassar A, Pellacani G, et al. In Vivo and Ex Vivo Confocal Microscopy for Dermatologic and Mohs Surgeons. Dermatol Clin. 2016;34(4):497–504.

2. van Royen ME, Verhoef EI, Kweldam CF, van Cappellen WA, Kremers GJ, Houtsmuller AB, et al. Three dimensional microscopic analysis of clinical prostate specimens. Histopathology. 2016;69(6):985–92.

3. Cahill LC, Fujimoto JG, Giacomelli MG, Yoshitake T, Wu Y, Lin DI, et al. Comparing histologic evaluation of prostate tissue using nonlinear microscopy and paraffin H&E: a pilot study. Mod Pathol. 2019 Jul 1;32(8):1158–67.

4. Tao YK, Shen D, Sheikine Y, Ahsen OO, Wang HH, Schmolze DB, et al. Assessment of breast pathologies using nonlinear microscopy. Proc Natl Acad Sci U S A. 2014;111(43):15304–9.

5. Sun CK, Kao CT, Wei ML, Chia SH, Kärtner FX, Ivanov A, et al. Slide-free imaging of hematoxylin-eosin stained whole-mount tissues using combined third-harmonic generation and three-photon fluorescence microscopy. J Biophotonics. 2019;(September 2018):1–21.

6. You S, Tu H, Chaney EJ, Sun Y, Zhao Y, Bower AJ, et al. Intravital imaging by simultaneous label-free autofluorescence-multiharmonic microscopy. Nat Commun [Internet]. 2018;9(1). Available from: http://dx.doi.org/10.1038/s41467-018-04470-8

7. Fereidouni F, Harmany ZT, Tian M, Todd A, Kintner JA, McPherson JD, et al. Microscopy with ultraviolet surface excitation for rapid slide-free histology. Nat Biomed Eng. 2017 Dec 1;1(12):957–66.

8. Weisi Xie, Ye Chen, Yu Wang, Linpeng Wei, Chengbo Yin, Adam K. Glaser, Mark E. Fauver, Eric J. Seibel, Suzanne M. Dintzis, Joshua C. Vaughan, Nicholas P. Reder, Jonathan T. C. Liu, “Microscopy with ultraviolet surface excitation for wide-area pathology of breast surgical margins,” J. Biomed. Opt. 24(2), 026501 (2019), doi: 10.1117/1.JBO.24.2.026501.

9. Yoshitake T, Giacomelli MG, Quintana LM, Vardeh H, Cahill LC, Faulkner-Jones BE, et al. Rapid histopathological imaging of skin and breast cancer surgical specimens using immersion microscopy with ultraviolet surface excitation. Sci Rep [Internet]. 2018;8(1):1–12. Available from: http://dx.doi.org/10.1038/s41598-018-22264-2

10. Wang M, Tulman DB, Sholl AB, Kimbrell HZ, Mandava SH, Elfer KN, et al. Gigapixel surface imaging of radical prostatectomy specimens for comprehensive detection of cancer-positive surgical margins using structured illumination microscopy. Sci Rep [Internet]. 2016;6(May):1–16. Available from: http://dx.doi.org/10.1038/srep27419

11. Wang M, Kimbrell HZ, Sholl AB, Tulman DB, Elfer KN, Schlichenmeyer TC, et al. High-resolution rapid diagnostic imaging of whole prostate biopsies using video-rate fluorescence structured illumination microscopy. Cancer Res. 2015;75(19):4032–41.

12. Glaser AK, Reder NP, Chen Y, Mccarty EF, Yin C, Wei L, et al. Light-sheet microscopy for slide-free non-destructive pathology of large clinical specimens. Nat Biomed Eng. 2018;1(7):1–22.

13. Reder NP, Glaser AK, McCarty EF, Chen Y, True LD, Liu JTC. Open-Top Light-Sheet Microscopy Image Atlas of Prostate Core Needle Biopsies. Arch Pathol Lab Med. 2019;143(9):1069–75.

14. Glaser AK, Reder NP, Chen Y, Yin C, Wei L, Kang S, et al. Multi-immersion open-top light-sheet microscope for high-throughput imaging of cleared tissues. Nat Commun [Internet]. 2019;10(1):1–8. Available from: http://dx.doi.org/10.1038/s41467-019-10534-0

15. Rocha MD, Düring DN, Bethge P, Voigt FF, Hildebrand S, Helmchen F, et al. Tissue clearing and light sheet microscopy: Imaging the unsectioned adult zebra finch brain at cellular resolution. Front Neuroanat. 2019;13(February):1–7.

16. Gareau DS, Li Y, Huang B, Eastman Z, Nehal KS, Rajadhyaksha M. Confocal mosaicing microscopy in Mohs skin excisions: feasibility of rapid surgical pathology. J Biomed Opt. 2008;13(5):054001.

17. Bini J, Spain J, Nehal K, Hazelwood V, DiMarzio C, Rajadhyaksha M. Confocal mosaicing microscopy of human skin ex vivo: spectral analysis for digital staining to simulate histology-like appearance. J Biomed Opt. 2011;16(7):076008.

18. Giacomelli MG, Husvogt L, Vardeh H, Faulkner-Jones BE, Hornegger J, Connolly JL, et al. Virtual hematoxylin and eosin transillumination microscopy using epi-fluorescence imaging. PLoS One. 2016;

19. Macenko M, Niethammer M, Marron JS, Borland D, Woosley JT, Guan X, et al. A method for normalizing histology slides for quantitative analysis. Proc - 2009 IEEE Int Symp Biomed Imaging From Nano to Macro, ISBI 2009. 2009;1107–10.

20. Rana A, Yauney G, Lowe A, Shah P. Computational Histological Staining and Destaining of Prostate Core Biopsy RGB Images with Generative Adversarial Neural Networks. Proc - 17th IEEE Int Conf Mach Learn Appl ICMLA 2018. 2019;828–34.

21. Rivenson Y, Wang H, Wei Z, Haan K De, Zhang Y, Wu Y, et al. Virtual histological staining of unlabelled tissue-autofluorescence images via deep learning. Nat Biomed Eng [Internet]. 2019;3(June). Available from: http://dx.doi.org/10.1038/s41551-019-0362-y

22. Rizwan A, Bulte C, Kalaichelvan A, Cheng M, Krishnamachary B, Bhujwalla ZM, et al. Metastatic breast cancer cells in lymph nodes increase nodal collagen density. Sci Rep. 2015 May 7;5.

23. Borhani N, Bower AJ, Boppart SA, Psaltis D. Digital staining through the application of deep neural networks to multi-modal multi-photon microscopy. Biomed Opt Express. 2019;10(3):1339.

24. Rivenson Y, Liu T, Wei Z, Zhang Y, de Haan K, Ozcan A. PhaseStain: the digital staining of label-free quantitative phase microscopy images using deep learning. Light Sci Appl [Internet]. 2019;8(1). Available from: http://dx.doi.org/10.1038/s41377-019-0129-y

25. Rudin C. Stop explaining black box machine learning models for high stakes decisions and use interpretable models instead. Nat Mach Intell [Internet]. 2019;1(5):206–15. Available from: http://dx.doi.org/10.1038/s42256-019-0048-x

26. Lam SK, Pitrou A, Seibert S. Numba: A LLVM-based python JIT compiler. Proc Second Work LLVM Compil Infrastruct HPC - LLVM ‘15. 2015;1–6.

27. Hörl D, Rojas Rusak F, Preusser F, Tillberg P, Randel N, Chhetri RK, et al. BigStitcher: reconstructing high-resolution image datasets of cleared and expanded samples. Nat Methods. 2019;16(9):870–4.

28. Gutman DA, Khalilia M, Lee S, Nalisnik M, Mullen Z, Beezley J, et al. The digital slide archive: A software platform for management, integration, and analysis of histology for cancer research. Cancer Res. 2017;77(21):e75–8.

29. Ruifrok, A. C. & Johnston, D. A. Quantification of histochemical staining by color deconvolution. Anal. Quant. Cytol. Histol. 23, 291–299 (2001).

30. Van Der Walt S, Schönberger JL, Nunez-Iglesias J, Boulogne F, Warner JD, Yager N, et al. Scikit-image: Image processing in python. PeerJ. 2014;2014(1):1–18.

31. Otsu N. THRESHOLD SELECTION METHOD FROM GRAY-LEVEL HISTOGRAMS. IEEE Trans Syst Man Cybern. 1979;

32. Anaconda Software Distribution. cudatoolkit version: 10.2.89 anaconda.org/anaconda/cudatoolkit

